# DriverSEAT: A spatially-explicit stochastic modelling framework for the evaluation of gene drives in novel target species

**DOI:** 10.1101/2022.06.13.496025

**Authors:** Mathieu Legros, Luke G. Barrett

## Abstract

Gene drives represent a potentially ground breaking technology for the control of undesirable species or the introduction of desirable traits in wild population, and there is strong interest in applying these technologies to a wide range of species across many domains including agriculture, health, conservation and biosecurity. There remains however considerable uncertainty regarding the feasibility and efficacy of gene drives in various species, based in particular on biological and ecological specificities of each target. In this paper we introduce DriverSEAT, a new spatial, modular modelling framework designed to assess the outcome of gene drives in a range of target species based on their specific ecological dynamics and genetics. In addition to the main structure and characteristics of the model, we present an example of its application on scenarios of genetic control of weeds, a potential candidate for gene drive control that presents significant challenges associated with plant population dynamics. We illustrate here how the results from DriverSEAT can inform on the potential value of gene drives in this specific context, and generally provide ecologically informed guidance for the development and feasibility of gene drives as a control method in new target species.

## Introduction

Gene drive approaches to the control of undesirable species have attracted a lot of interest and research over the past decade or so. Thanks in large part to the advent of novel gene editing technologies based around the use of the CRISPR-Cas family of site-directed nucleases, the feasibility of these techniques has significantly improved in recent years, and they are now seen as a potential control tool against a wide range of target species. In practice, gene drives have successfully been engineered in several species, including insect species (Buchman et al., 2018; Gantz et al., 2015; Hammond et al., 2016; Li et al., 2020), rodents (Grunwald et al., 2019; Pfitzner et al., 2020) and fungal plant pathogens (Gardiner et al., 2020), and have proven to be capable of drive (i.e. an increase in frequency of the gene drive element beyond what Mendelian genetics would predict) in controlled laboratory settings.

Initial research efforts were primarily focused on targeting insect vectors that carry diseases that significantly impact public health, as well as invasive species that decimate island ecosystems (Godwin et al., 2019). Because of the success of these early proof of concepts, as well as the inherent versatility of the CRISPR tools used to engineer most of these constructs, there is now tremendous interest in the potential applications of these technologies to tackle problems in human health, agriculture and conservation.

In particular, gene drives are now being considered as potential tools for the control of many pests of agriculture (Legros et al., 2021) and natural environments (Godwin et al., 2019). There are many incentives that support the development of novel technologies for agricultural pest control. First and foremost are the significant crop losses and associated economic damage caused worldwide by a wide range of pest species. Second, while there are a variety of chemical, genetic and cultural methods currently used for pest control, the rapid evolution of pests against these control measures (e.g. host resistance, chemical pesticides) are of significant concern. Significant decreases in efficacy have been reported for first-line herbicides, fungicides and insecticides (Gould et al., 2018a). Novel control methods that reduce the reliance on chemical biocides and other control tools would likely represent a significant improvement in the durability and sustainability of pest control methods in many agricultural systems (Kumaran et al., 2020).

While the interest is therefore present, there are still many areas of uncertainty regarding the feasibility and potential suitability of gene drive approaches for the control of new target pests. The first hurdles are primarily related to the engineering itself of gene drive constructs in novel target species. As an example, potential applications in plants are likely to be faced with several specific difficulties (Barrett et al., 2019). In the present article, however, we focus on the uncertainties that relate more specifically to the fate of an existing gene drive when released into a target population and environment. In other words, we assume that a gene drive can be engineered, and address the questions regarding its potential efficacy as a pest control method.

Theoretical tools to assess the feasibility and potential outcome of a gene drive control strategies are a crucial part of the gene drive research pipeline, and several studies have focused on such theoretical evaluations (NASEM, 2016) (see (Golnar et al., 2021) for a recent review). Generic models of gene drive typically focus on the dynamics of a gene drive element in a simplified population (or metapopulation)(Backus and Delborne, 2019; Davis et al., 2001; Edgington and Alphey, 2018; Esvelt et al., 2014; Noble et al., 2018, 2019; Ward et al., 2011), and provide crucial insight on the overall fate of a hypothetical gene drive, the role of factors such as fitness costs (Backus and Delborne, 2019), population structure (Bull et al., 2019)(Huang et al., 2009, 2011) and resistance evolution (Unckless et al., 2017). In the context of evaluating gene drives as a control tool against specific target species, there is a need for more complex models that account for the biological and ecological specificities of the target, of its environment and its target landscape (Golnar et al., 2021). Such models have been developed for specific instances of gene drive control, primarily in mosquitoes, with models such as Skeeter Buster (Legros et al., 2012, 2013; Magori et al., 2009) or MGDrive (Sánchez C. et al., 2020; Wu et al., 2021). However, these models are designed to specifically simulate mosquito populations (or, in the case of the former, a single mosquito species), and therefore lack the versatility required for a comprehensive gene drive evaluation tool at a time where potential target species cover a wide range of taxa.

To fill that gap in the current collection of available theoretical tools, we developed and present here DriverSEAT (multi-Species Ecological Assessment Tool), a modelling framework designed to study the dynamics of a gene drive in biologically and ecologically detailed, spatially structured simulated populations. Because we aim to investigate the feasibility of gene drive methods when applied to a variety of pest species, we developed a generic and modular theoretical framework that can be tailored and customised to accommodate a wide range of target species, population structures and environmental conditions. In the current article we introduce the overall structure of DriverSEAT, its main components and its potential applications. In addition, as an illustration of the applications of this modelling framework, we present a specific investigation of potential gene drive applications for the control of agriculturally significant weeds. To this day there has been only one example of a functioning gene drive engineered in plants, namely *Arabidopsis thaliana* (Zhang et al., 2021). There consequently remains considerable uncertainty regarding the feasibility of such control methods (Neve, 2018). We therefore choose the example of gene drive in plants to illustrate how our modelling framework addresses the uncertainties related to gene drive development in novel target species. In particular we focus on the impact of traits relevant for potential target weed species: selfing, seed banks and spatial heterogeneity. We show how each of these aspects of the biology of a target species can help or hinder the spread of a gene drive (and conversely, render its confinement more or less challenging). We conclude by detailing the potential use for this modelling framework in future investigations of gene drives and their suitability as control methods for a variety of pests and target environments, where gene drives might constitute a promising addition to the collection of available control methods.

## Model description

In this section we detail the structure and parameterisation of our modelling framework. Because DriverSEAT is designed to accommodate a wide range of simulated species and scenarios, the precise functions used for most biological and ecological processes can vary and be specified according to need. In this section we aim to present the general characteristics of the framework and, where applicable, provide examples of specific procedures and mathematical functions, pertaining in particular to the case of gene drive applications in weeds, our example of choice in this article.

### Modelling framework

The core structure of DriverSEAT consists of a spatial, stochastic meta-population built as a collection of individual patches connected by links representing specific distances (see Spatial structure). Note that the spatial scale itself is not explicitly defined in this model, therefore each patch might represent a paddock, a farm, a county or a region, depending on the specific scale of interest. Within each patch, however, there is no further spatial structure, and within-patch populations are assumed to be spatially well mixed.

The model operates on discrete time, and the dynamics within patches and among patches are computed and calculated at every time step. A critical feature of this model is modularity. This tool was designed with the aim to accommodate the widest range of scenarios possible, both in terms of target species and environments. At the same time, it was developed to operate only with the parameters and functions that are directly relevant to the considered scenario. Therefore, while we list in this section a lot of features that can be included in runs of the model, it should be noted that not all features are required for a given application of the model and can be conveniently turned off.

Within each patch the dynamics of two distinct species can be independently modelled. The dynamics of each species can be described by its own equation, either independently with e.g. logistic growth or periodic dynamics driven by environmental factors, or as a system of equations describing a range of biotic interactions, including competition, predator-prey or host-parasite interactions. Here again, the modularity of the system ensures that a variety of biological and environmental settings (like a fungal pathogen and its host plant, a resident species and its invasive competitor, or an insect pest and its natural enemy) can be easily simulated within this particular framework.

### Spatial structure and population dynamics

The fundamental spatial structure of the model consists of a collection of individual sites organized in a network with defined coordinates for each site and distances between each site. In this article, this metapopulation is organized as a square lattice with contiguous sites, so that the distance between two given sites is equal to the Euclidean distance between them in the coordinate system defined by the lattice. Boundary conditions are set to be periodic (opposite edges of the lattice are connected) so that the population can be considered as one toroidal spatial unit with no boundaries.

The population dynamics within each site are tracked following a discrete numerical implementation of the model chosen to describe the within-cell dynamics of the species of interest. In the simplest case, this will be a one-compartment model with logistic density-dependent growth within each cell. Our modular framework can, however, accommodate a variety of more elaborate population dynamics models within each site. This allows in particular to take into account more elaborate age- and stage-structured dynamics. In this paper we present one such example applied to plant populations (see next section) though other types of age-structured dynamics can be implemented.

Note that there is no inherent size associated with any individual model cell, and no specific unit for the distance between cells. Therefore, DriverSEAT could be conceptually applied to a variety of spatial scales, whereby an individual cell in the model could represent a plant in a paddock, a paddock in a farm, a farm in a landscape or any administrative subdivision at even larger scales. In our example in this article, we assume that each cell represents an individual paddock, and that the model grid as a whole constitutes an abstract representation of an agricultural landscape.

### Life history traits - Dispersal

Dispersal in the most general sense in this model is a function that ensures the movement of propagules from one time step to the next, from one patch to another patch. Depending on the target species, the propagules might be seeds, pollen, spores, migrating organisms, or any other dispersing element. In the model this process is represented as a two-step process: one defines the probability of dispersal occurring from a given patch at a given time, the other describes the dispersal kernel from which the respective probabilities of any other site being chosen as the destination site are calculated.

This dispersal kernel is defined as a probability density function that described dispersal probability as a function of distance. The model can accept essentially any mathematical function as a dispersal kernel, although it is important that the user ensures that the dispersal distances are correctly set in relation to the distances between sites defined as the metapopulation spatial structure.

Dispersal patterns can be extraordinarily diverse across taxa and species, and the choice of mathematical functions to describe them is virtually infinite. In the context of this paper, where we use weedy plants as an example target species, we choose to use a dispersal kernel for seeds and pollen that describes an anisotropic wind-mediated dispersal (see next section) as the most common dispersal mechanisms among plant species.

### Genetics

Genotypes in the model are represented by a bit string where each locus is encoded by two bits, thereby allowing up to 4 alleles. As an example, a genotype represented by the following bit string:

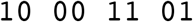

represents an individual genome with 4 loci, with allele ‘01’ at the first (rightmost) locus, allele ‘11’ at the next locus, etc. This genotype can be stored as the integral representation of this bitstring, i.e. in decimal form, 141.

Genotypes are stored as an `unsigned long long int` type in the model, with a capacity of 64 bits. The model can therefore accommodate up to 32 loci in haploid organisms, 16 loci in diploid organisms, and ⌊32/*m*⌋ loci in *m*-ploid organisms.

Each locus can be associated with its own inheritance pattern, allowing for various types of gene drives to be described simultaneously in the model (as well as non-drive genetic elements, including selective and neutral elements). Each gene drive is defined by the set of probabilities that an offspring with genotype *g* results from the mating of parents with genotypes *I* and *j* (need to explicit those probabilities for various gene drives?)

The genetic composition of the population in each model cell is stored as an array containing the proportion of each genotype in the cell.

### Mating and sexual reproduction

At specified time steps, reproduction can occur within any site in the model’s metapopulation. The timing of reproduction events and proportion of reproducing individuals can be defined by the user and can be specified to occur (or not) at any time step. Reproducing individuals are chosen by randomly sampling from the existing genotypic composition within a particular site. Depending on the mechanism of reproduction for the species of interest, several options are then possible.

For asexual (clonal) reproduction, genotypically identical offspring are added to the same site, in proportion to the frequency of the parent of a specific genotype and their (genotype-specific) fecundity.

For sexual reproduction, the procedure depends on the mating patterns of the species of interest. A proportion *c* of selfing individuals can be defined, in which case each reproducing individual is designated to mate with itself with probability *c*. Otherwise a mate with suitable mating type is chosen at random within the same site.

Mating types are encoded in our model with each individual’s genotype. In the typical cases with only 2 mating types (2 sexes), one locus in the genotype encodes the individual’s sex, 00 for females and 01 for males. However there is considerable flexibility within this structure to encode for more mating types and/or more mating loci, a likely relevant factor for fungal populations (Billiard et al., 2012).

### Model implementation

DriverSEAT was developed in C++ and uses the GNU Scientific Libraries publicly available at http://www.gnu.org/software/gsl/. The source code is available at https://github.com/legrosmathieu/. Documentation for the model is available in the same repository.

## Parameterisation and simulations

### Case study: genetic control of weed populations

Weed species cause significant ecological and economic damage to many environments across the world. Weeds are a particular concern to agriculture, and the associated economic costs, while difficult to estimate, are very substantial. For example, annual costs of weed damage and control has been estimated at AUD 3.3 billion in Australian grain production (Llewellyn et al., 2016), costs associated with crop and pasture weeds in the USA were evaluated at over USD 32 billion annually (Pimentel et al., 2000), and overall annual costs to agricultural production in India were estimated in excess of USD 11 billion (Gharde et al., 2018). In addition to their economic significance, weeds also represent an increasingly challenging target for control, in large part due to the emergence and spread of resistance to herbicides (Gould et al., 2018b).

Given this situation, the idea of gene drives as a new tool for the control of weed populations has generated a lot of interest. There are however many significant challenges to the implementation of such a control option, including obstacles specifically linked to the engineering of gene drives in plant species (Barrett et al., 2019; Kumaran et al., 2020), as well as challenges associated with genetic control in agricultural environments (Legros et al., 2021). With so many potential hurdles, and therefore such levels of uncertainty, theoretical tools are especially valuable for assessing the feasibility of gene drives as a weed control option as an early step in the R&D process. We have therefore chosen to investigate the hypothetical control of an agriculturally significant weed, and describe here how we tailor our modelling framework to this specific case study.

### Weed population structure: seed banks

Age (or stage) structure plays a significant role in the outcome of a gene drive (Huang et al., 2009). For plants, we identified seed banks as the component of age structure most likely to impact gene drive, as suggested by earlier studies of population genetics and evolution within seed banks (Koopmann et al., 2017; Shoemaker and Lennon, 2018). We show here the example of a compartmental model that described the dynamics of an annual plant with a persistent seed bank across seasons (see Supplementary Figure 1). The pattern of seed dormancy across seasons is followed by tracking seed of age *a* and genotype *g* in specific compartments *S_g,a_*. The transitions between compartments are defined by age- and genotype-specific viability *μ_g,a_* and germination rates *b_g,a_*. The above ground population is tracked for each genotype *g* without any additional age structure. These above ground individuals are reproducing each year (for an annual plant) and contribute new seeds to the *S_g,0_* compartments.

### Weed dispersal mechanisms

For weed populations, we choose to model the most common propagule (pollen and seeds) dispersal mechanism, namely wind-mediated dispersal. In mathematical terms, to describe an anisotropic wind-mediated dispersal kernel, we chose an exponential-power distribution, that has been shown to best describe this type of dispersal (Bullock et al., 2017):

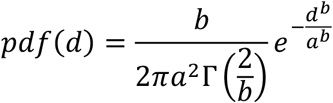

where *d* is the distance between sites, and Γ is the gamma function:

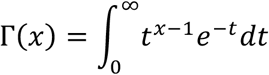

The dispersal kernel is described by two parameters: a_s_ and b_s_ for seeds (respectively a_p_ and b_p_ for pollen). The impact of each parameter on the shape of the kernel is described in Supplementary Figure S2.

## Results

### Impact of life history traits and age structure on gene drive dynamics

The modularity of the DriverSEAT framework means that the model can be used to investigate the impact of a range of key traits, specific to a target of interest, on the outcome of a gene drive strategy. As an illustration, we use the example of an annual weed species with a Site-Directed Nuclease (SDN)-type drive, and focus on two key biological traits: inbreeding and seed banks. While these two elements have been shown in more general models to impact drive dynamics in plant populations (Barrett et al., 2019; Bull et al., 2019), here we examine their respective importance in a more quantitative fashion.

As expected, an increased selfing rate significantly hinders the spread of a gene drive in a theoretical weed population, as does the error rate in the SDN drive mechanism (where *e* is the probability of generating a resistant allele through error-prone repair mechanisms like non-homologous end joining). For example, for a hypothetical error-free SDN drive (*e*=0), the frequency of the drive allele reaches 90% after 10 generations in a purely outcrossing population (*F*=0), but only reaches 50% in the same time frame with a moderate selfing rate (*F*=0.2) (Fig 1). Overall, selfing has a substantial impact on the fate of the gene drive element even at relatively low rates (no higher than 20% in this example). We also note that the impact is strongest on the gene drives with low rates of resistant allele generation, illustrating that, in this scenario, selfing and resistance can interact to negate the successful spread of a gene drive element.

**Figure 1.**
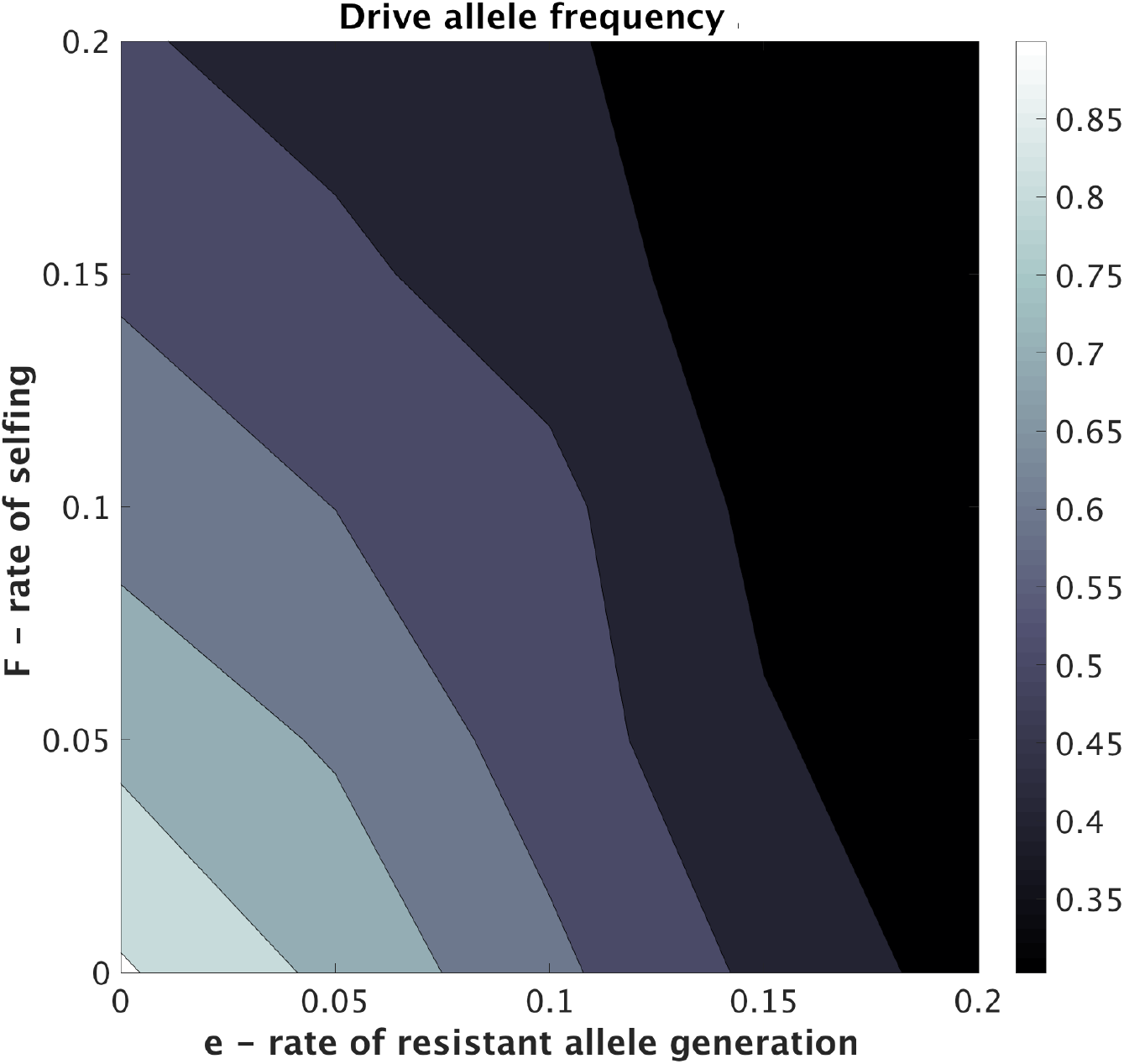
Impact of selfing rate on the outcome of a SDN gene drive in a plant population after 10 generations.

We then illustrate the consequences of introducing a seed bank in the dynamics of this hypothetical annual weed species. The basic structure of the seed bank model is illustrated in Supp. Figure 1. We simulate the release of an SDN drive carrying a recessive lethal genetic cargo as a population suppression strategy (Figure 2). Compared to a scenario with no seed bank (ν=1.0), where the gene drive reaches 90% frequency after 9 generations, the existence of a seed bank (ν<1.0) substantially slows down the spread of the drive, and consequently delays the population suppression effect imposed by the recessive lethal cargo. A 95% population reduction can be achieved in 11 generations in a species without a seed bank, whereas the same level of suppression would take over 50 generations in a species with a long-lived seed bank (ν=0.2, corresponding to an average time of 5 years spent in the seed bank). While the former might be considered a viable long-term control solution for a problematic annual weed, the latter in most cases would likely be an unacceptable time frame.

**Figure 2.**
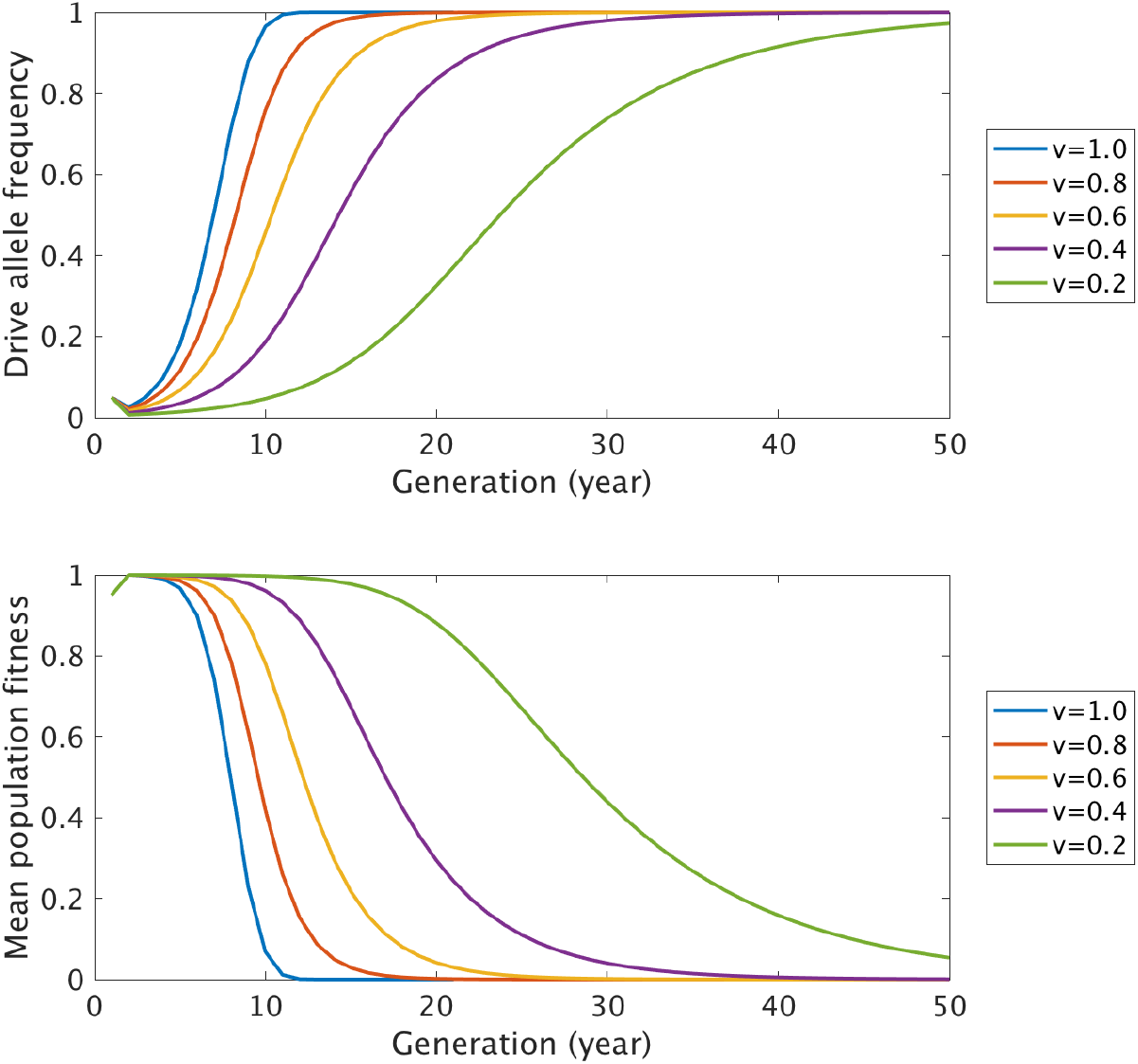
Impact of the presence of a seed bank and of the average time to germination on the outcome of a population suppression strategy based on the release of an SDN drive carrying a recessive lethal genetic cargo.

### Gene flow and gene drive spread in a weed population

Dispersal is a crucial element for the spread of a gene drive amongst and between populations (including for its potential spread to non-target areas or populations). This is true in plants as in any other target, and for our example of an annual weed species, we choose to investigate the impact of the dispersal patterns of both pollen and seeds on the outcome of an SDN gene drive.

We use isotropic, wind-mediated patterns to describe the dispersal kernels for both types of propagules. As described above, we selected an exponential-power distribution as its mathematical description (Bullock et al., 2017). With this function we can modify two parameters (*a* and *b*) to impact the scale (*a*) and shape (*b*) of the dispersal kernel (Supp. Figure 2). In this context we show that the fate of a gene drive, when starting from low frequencies in a population of an annual weed species, is most strongly impacted by the average dispersal distance of pollen *(a_pollen_)* especially for *a_pollen_*<=2 (Figure 3). We observe that the average dispersal distance of seeds *(a_seed_)* also has a substantial impact on the fate of the gene drive, albeit quantitatively less important than for pollen. The shape of the respective kernels *(b_pollen_* and *b_seed_)* appears to have little quantitative impact on the final frequency of a gene drive, but significantly increase the variability observed among replicated runs in our model, suggesting that more frequent long distance dispersal events have a destabilising effect on the ability of a low frequency gene drive to spread in a local population.

**Figure 3.**
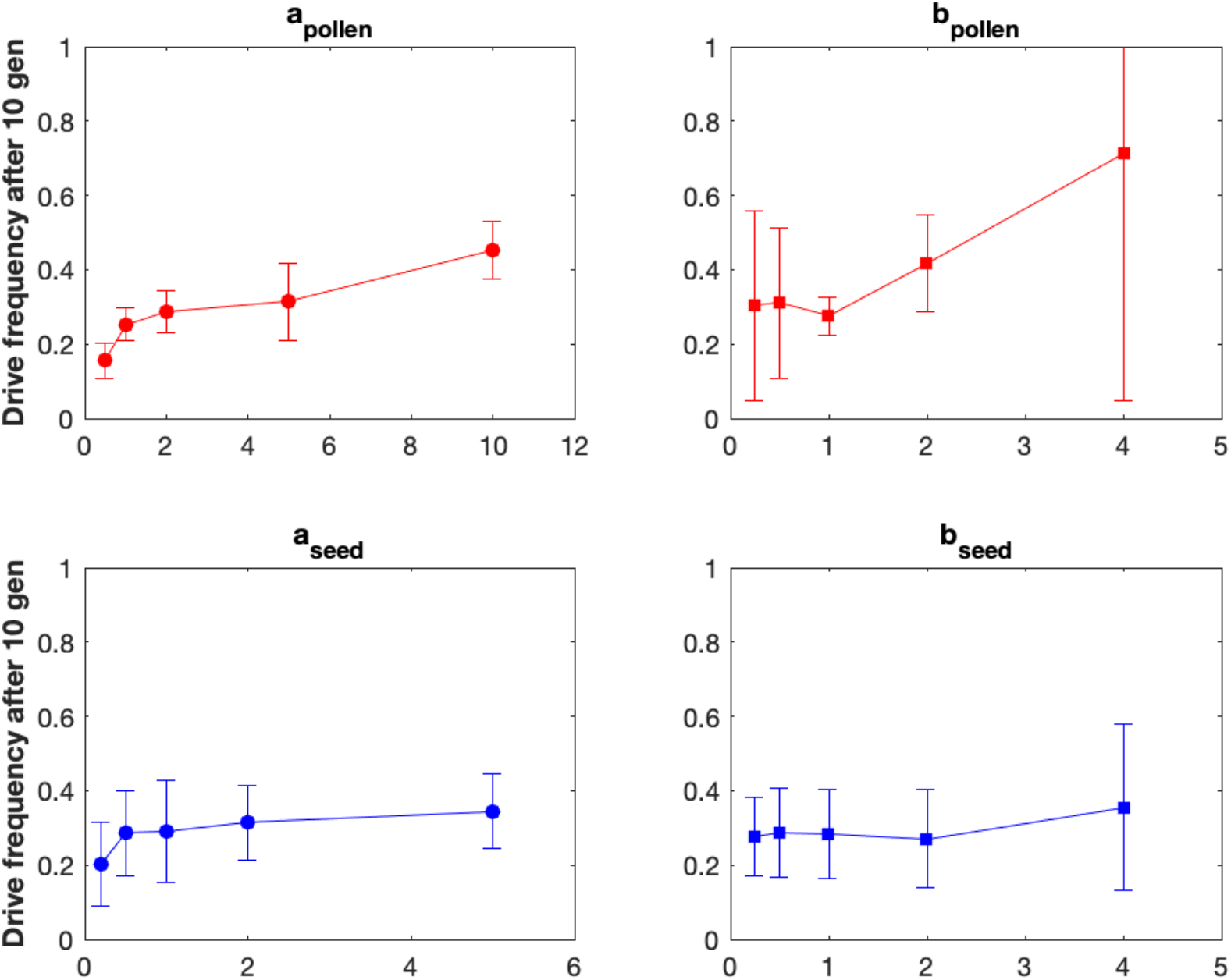
Dispersal kernels for pollen and seeds and outcome of a gene drive in an annual weed

### Spatial dynamics and deployment

At the population level, determining the strategy for release of gene drives into target environments (e.g. agricultural paddocks within landscapes) presents another challenge, the precise nature of which depends on the species of interest and the chosen genetic control approach. To illustrate how our modelling framework can inform decisions around spatial dynamics and deployment, we again simulate the case of an annual weed in an agricultural landscape, and the release of an SDN-based gene drive designed to drive a trait of interest into the population (e.g. herbicide sensitivity).

Figure 4 depicts the dynamics of such a gene drive following its introduction into a single site. The model predictions show that this non-localized drive (i.e. with potential to spread throughout the landscape) unsurprisingly spreads easily across the metapopulation in the model. More importantly, these simulations can be used to understand the dynamics of spread within the landscape. For instance, we can make quantitative predictions regarding the expected delay before a gene drive invades and spreads into a patch depending on distance from a release site. While such distances in the model are expressed in arbitrary units, it would be easy to translate this information into actual distances depending on mean patch size within the environment of interest. Whether the spread of such a gene drive is seen as a desired outcome (for area-wide control) or an unintended consequence (spread into non-target areas) will depend on the context that is being simulated. Nevertheless, the ability to make such quantitative predictions and risk evaluations will be valuable in either case.

**Figure 4.**
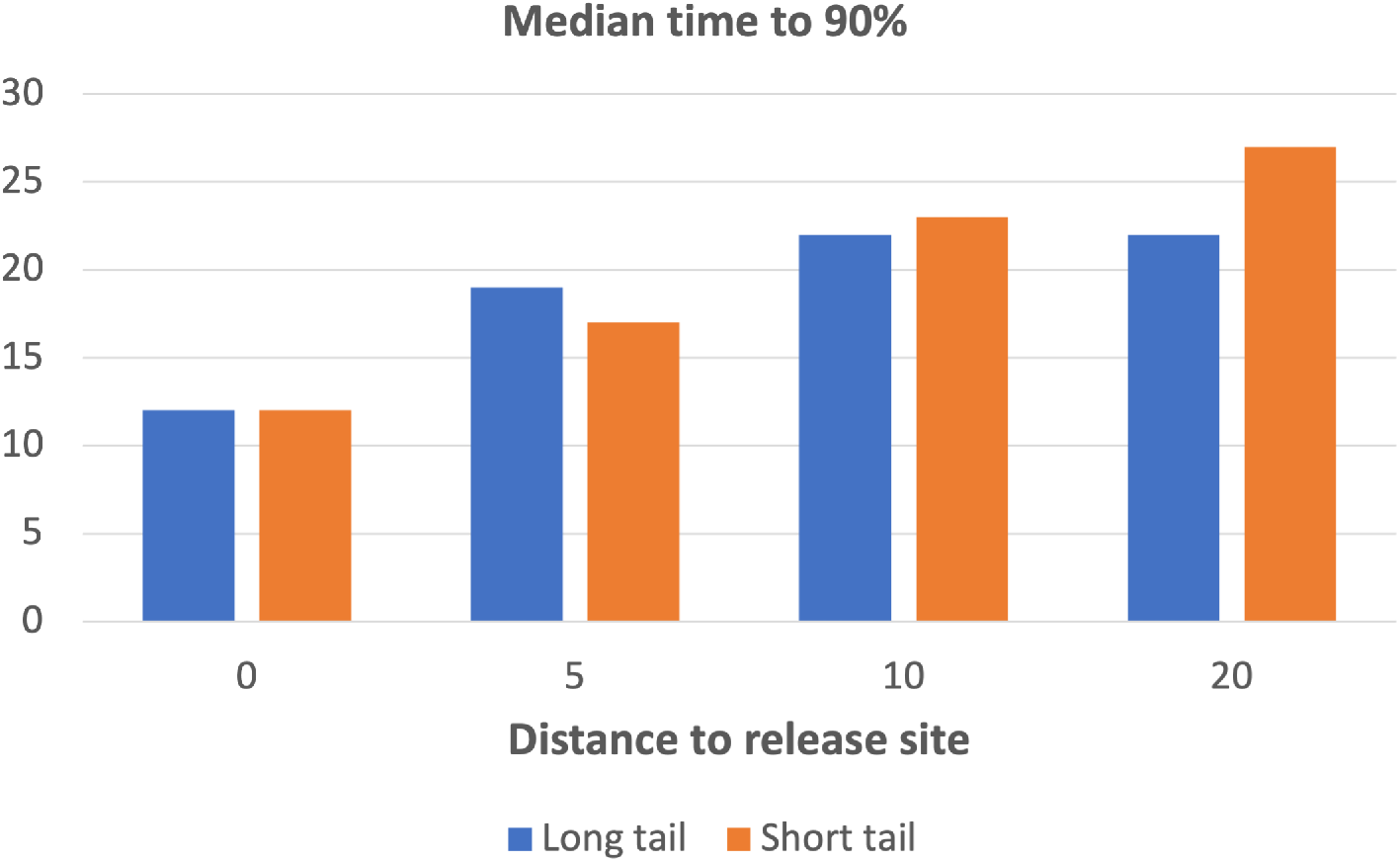
Median time to 90% frequency for an SDN-based gene drive after a single release in a single release site in the centre of a population. Long tail (b=1) vs short tail (b=3)

DriverSEAT was also designed to inform decisions regarding the optimal methods to deploy a gene drive into a target area. For a single release of an SDN-based gene drive, we can compare strategies of uniform release into every cell in the meta-population (potentially logistically challenging) to strategies of selected release sites (Figure 5). In this example we select a number of release areas (1, 4, 9 or 16) that are regularly chosen across the grid (respectively one central area, or a 2×2, 3×3 or 4×4 grid of release sites), while keeping the number of cells with a release and the number of released individuals constant. Interestingly we show (Figure 5) that with a carefully chosen regular grid of 16 release sites across a 20×20 overall population, the gene drive can spread at least as quickly across the entire area compared to a uniform release across the entire population. In more complex landscapes this modelling framework could therefore provide valuable guidance about the optimal patterns for the deployment of gene drives.

**Figure 5.**
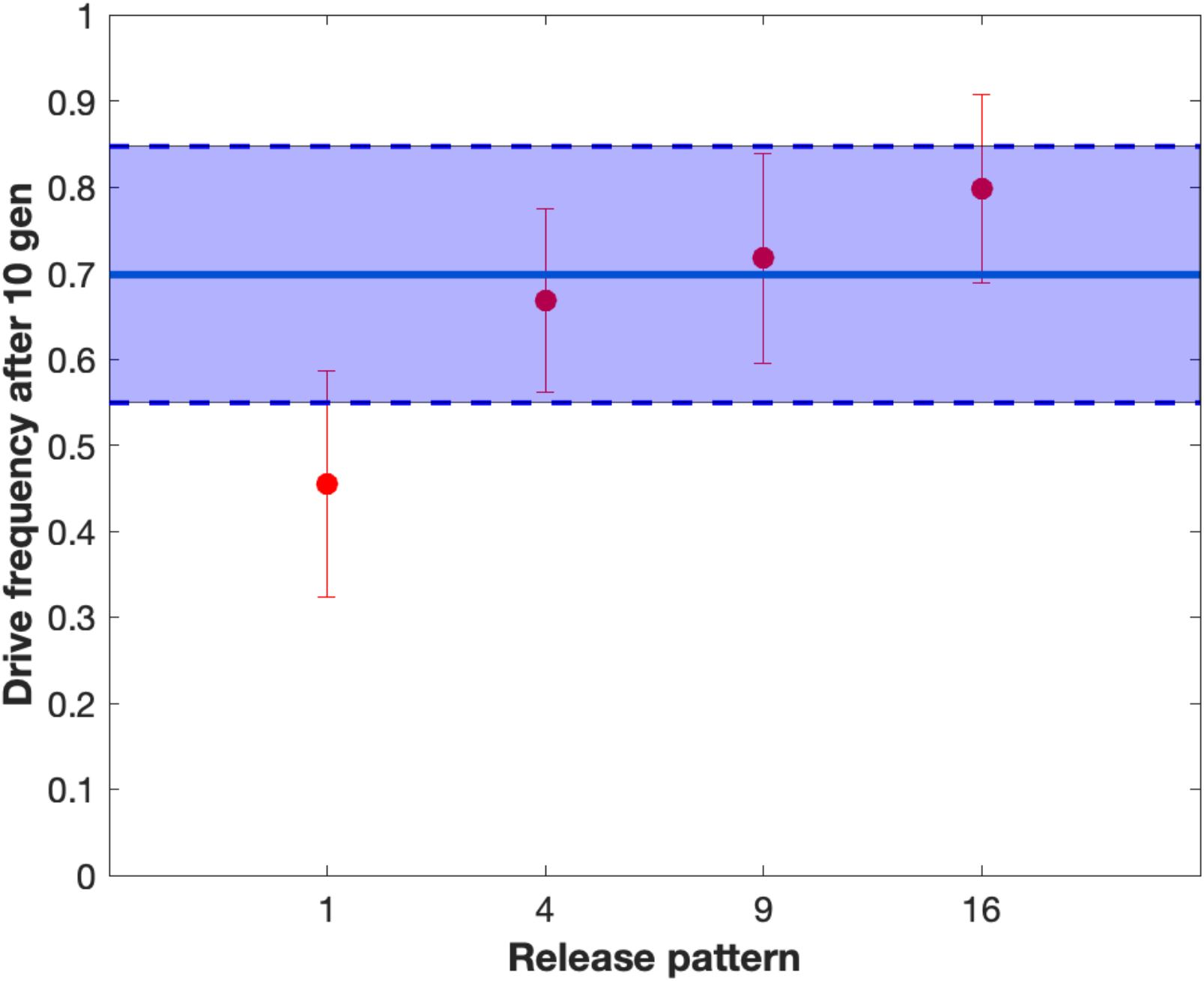
Impact of the release pattern on the outcome of SDN-based gene drive after a single release in the population. The value on the x-axis indicates the number of release sites selected in the population (note that the overall number of released individuals remains constant). The blue shaded area denotes the results of a uniform release in every single site.

## Discussion

In this article we introduce DriverSEAT, a novel modelling tool designed to assist development and decision making around gene drive control opportunities for a wide range of new potential target species. This modelling framework is designed to be modular, versatile and biology rich, to ensure that key biological and ecological aspects of any given target species are taken into account when considering the potential impact of genetic control approaches. Gene drive strategies are being considered as control tools for a variety of targets and environments, from agriculture to conservation and public health, with potential target species including fungi, plants, insects and vertebrates (NASEM, 2016). For each of these possible applications, gene drive control constitutes a promising option, but very significant uncertainties remain regarding feasibility, durability and acceptability. These uncertainties are not static, but rather vary across the biological spectrum of potential target species. For these reasons, there is a significant need for a modelling framework that can generate predictions about the outcome of gene drive strategies for a wide range of species and environments, while simultaneously pinpointing the traits and features of each target that are most relevant to gene drive dynamics. In particular, identifying features or life-history traits that constitute an inherent obstacle, or where uncertainty has the most significant impact, is critical.

The main motivation behind the development of this tool was therefore to assist decision making around the development of gene drive strategies and technologies in new target species and environments. In such cases, the road from gene-drive conceptual design through to the large scale implementation of a genetic control program will be long, paved with significant uncertainty and involving considerable research investment. With this model we aim to provide a theoretical decision-support tool to assist throughout the research and development process. Understanding how trait variation can influence gene drive dynamics will help identify candidate species where gene drive is most likely to succeed, thereby reducing risks for research and development of a new technology. The model can also be used to identify optimal approaches in a particular context (e.g. high-threshold vs. non-localized drives), or guide experimental research and data collection to most efficiently reduce uncertainty about the outcome of a particular gene drive strategy.

Gene drives offer potential solutions for the management of a diverse range of pest organisms. Consequently, the model was developed with versatility and modularity as key features, and so can provide insight into drive dynamics and impact at different levels, from generic to the more species-specific. Here we have adopted a broad focus, illustrating the influence that key life-history traits might have when using gene drive in plants for weed control. Our results show how the model can provide insight into the influence of variation in traits that are generally relevant for many plant species, such as self-fertilization, seed banks and pollen and seed dispersal. While broad, such information can be used to provide specific guidance for the development of gene drive control methods in plants. For example, our results provide insight into the levels of inbreeding that are likely to be insurmountable for the successful use of SDN-based gene drives (Figure 1). While the relevance of our quantitative results will vary from species to species, we note that relatively low levels of selfing (up to 20%) seem to represent a significant hindrance to the spread of a gene drive. This is noteworthy because selfing rates exceeding 20% are commonly observed in many plant species (Charlesworth, 2006). Our results also emphasize the need for robust data collection around dispersal and population structure (especially for rare, long range dispersal events) so as to predict the long-term outcome of a gene drive release (Figure 5). For entirely novel targets like plants, where gene drives are in their infancy (only one example of an engineered gene drive in *Arabidopsis thaliana* has been published to date (Zhang et al., 2021), such high-level guidance can be used to focus research and development efforts towards viable species.

Ultimately DriverSEAT can be used to identify obstacles that may prove insurmountable for some candidate target species and therefore in turn guide the choice of optimal candidates for successful gene drive development and engineering. In the context of the examples simulated above, for our results demonstrate that gene drives have highest potential for success as a control method in outcrossing plant species with limited seed dormancy across seasons. Such information can provide valuable insight into decisions regarding the choice of an optimal target species, and help define what constitutes success and failure of a gene drive based approach to pest control.

Take, for example, a scenario involving development of gene drive strategies for the control of weeds of broad-acre grain farming in temperate Australia, where weeds cost the industry in excess of AU$3 billion annually (Llewellyn et al., 2016). Two highly problematic and challenging species that present as potential candidates for a gene drive strategy are wild radish (*Raphanus raphanistrum*) and annual rye grass (*Lolium rigidum*). These two species comprise a significant component of the overall economic impact on the Australian grains industry, with average calculated costs of AU$53 M and AU$93M per annum for wild radish and annual rye grass (Llewellyn et al., 2016). With common and widespread resistance emerging to registered herbicides, the management of these two weeds is an increasingly vexing problem. Thus, they present as good hypothetical candidates for the development of control strategies involving gene drive. Importantly, as the above results show, both species are effectively 100% outcrossing. However, one key life-history difference is that *R. raphanistrum* displays prolonged seed longevity in the soil compared to *L. rigidum* (Bajwa et al., 2021). As our general results above show, prolonged seed dormancy is likely to significantly delay the rate at which drive will occur. Thus, *prima facie*, it can be predicted that in the Australian context, *L. rigidum* is the better candidate for investment in gene drive development and deployment.

In any case, our modelling framework can then be tailored and calibrated to simulate the dynamics of target populations of the species of interest. This includes selecting the relevant modules to run within the model (e.g. here, age structure in the form of seed banks), eliminating irrelevant modules (e.g. inbreeding for obligate outcrossing species) and of course collecting data for species-specific parameterisation. This modelling framework can then provide quantitative assessment of the potential outcome of a variety of gene drive approaches in specific target populations of the species of interest. For instance, following up on the results presented in this article, we are currently investigating how this model can inform the development of gene drive approaches in the aforementioned weeds in Australian agricultural environments (i.e. annual ryegrass and wild radish). We show that this modelling framework can also be valuable at the time a hypothetical gene drive has been developed and is ready for release, particularly for the design and release strategies and patterns of engineered individuals. While we only discussed broad guidelines in this article, e.g. regarding uniform spatial distribution of releases (Figure 5), challenges related to release are likely to be very variable depending on the target species. In our example, uniform release patterns will likely be easier to achieve with the release of seeds for weed control, or eggs for mosquito control, but will likely prove significantly more challenging in the context of mobile animals (e.g mammal control programs).

Overall, DriverSEAT can provide evidence-based guidance for the development of gene drives in new targets at several different levels, from generic results about relevant biological and ecological features to specific quantitative assessment of the impact of any given trait or feature, or of the uncertainty around it, on the outcome of a gene drive control strategy. Thus, we anticipate that this modelling framework will constitute a valuable tool for any decision maker (e.g. scientist, regulator or stakeholder) involved in research and development around gene drives in novel target species. On the other hand, the model has limitations. Its versatility and wide range of potential target species means that it is likely not the ideal tool for precise simulations of a specific gene drive product in a specific target population (Legros et al., 2012, 2013; Xu et al., 2010), something that is likely to be of interest later in the gene drive research and development pipeline, close to large scale field testing. In such instances, where precise quantitative predictions about the fate of a gene drive in a specific area are needed, there will likely be a need for more specialized modelling tools, such as the ones that have been developed for current gene drive research programs in mosquitoes (Sánchez C. et al., 2020; Wu et al., 2021) and rodents (Prowse et al., 2019).

Additionally, in its current form our modelling framework does not include any detailed genomic information about the target population, and therefore cannot be used to assist in the development of features that are sensitive to intra-population genomic variation, e.g. the choice of target sites for SDN-based gene drives (Sudweeks et al., 2019). These important issues require the help of dedicated modelling tools, although, by specifying the rate at which resistance can emerge at the population level, our model can be used to inform on the impact of such variability in the long term.

Overall, this illustrates that for a long, often contentious and ambitious process like the development of novel, gene drive control technologies in a variety of new species, a wide range of such modelling tools will likely be needed, from the early conceptual research on gene drive feasibility to the late stage, field trials and associated uncertainty and risk assessments. In our opinion, DriverSEAT will be a valuable addition to this process, by helping decision making around the development of gene drive control strategies in new target species, including new taxa. In this article we have demonstrated our model can provide generic guidance regarding gene drives in plants, and where obstacles and uncertainties might be found. Thanks to its versatility, this model can provide further guidance in this context and beyond, both for more species-specific gene drive evaluations as well as for other candidate targets beyond weeds.

More generally, our results emphasize the importance of accounting for the biology and the ecology of the target species and environments when evaluating a potential gene drive strategy. Regardless of the specific setting, gene drives, if and when they are implemented, will always involve the release of organisms in a natural population, and genes persisting across generations (to varying degrees depending on the chosen genetic approach of course). Ecological and evolutionary dynamics will inevitably be at play and are likely to have a strong impact on the success or failure of a gene drive (or, perhaps just as importantly, the success of failure of its containment)(Marshall, 2009). It is crucial that these aspects be taken into consideration early on in the evaluation process for gene drives in new targets and new settings, making DriverSEAT, the modelling tool presented here, complementary to the collection of models that will accompany the development of these new, promising yet controversial technologies.

## Supplementary figures

**Supp. Figure 1.**
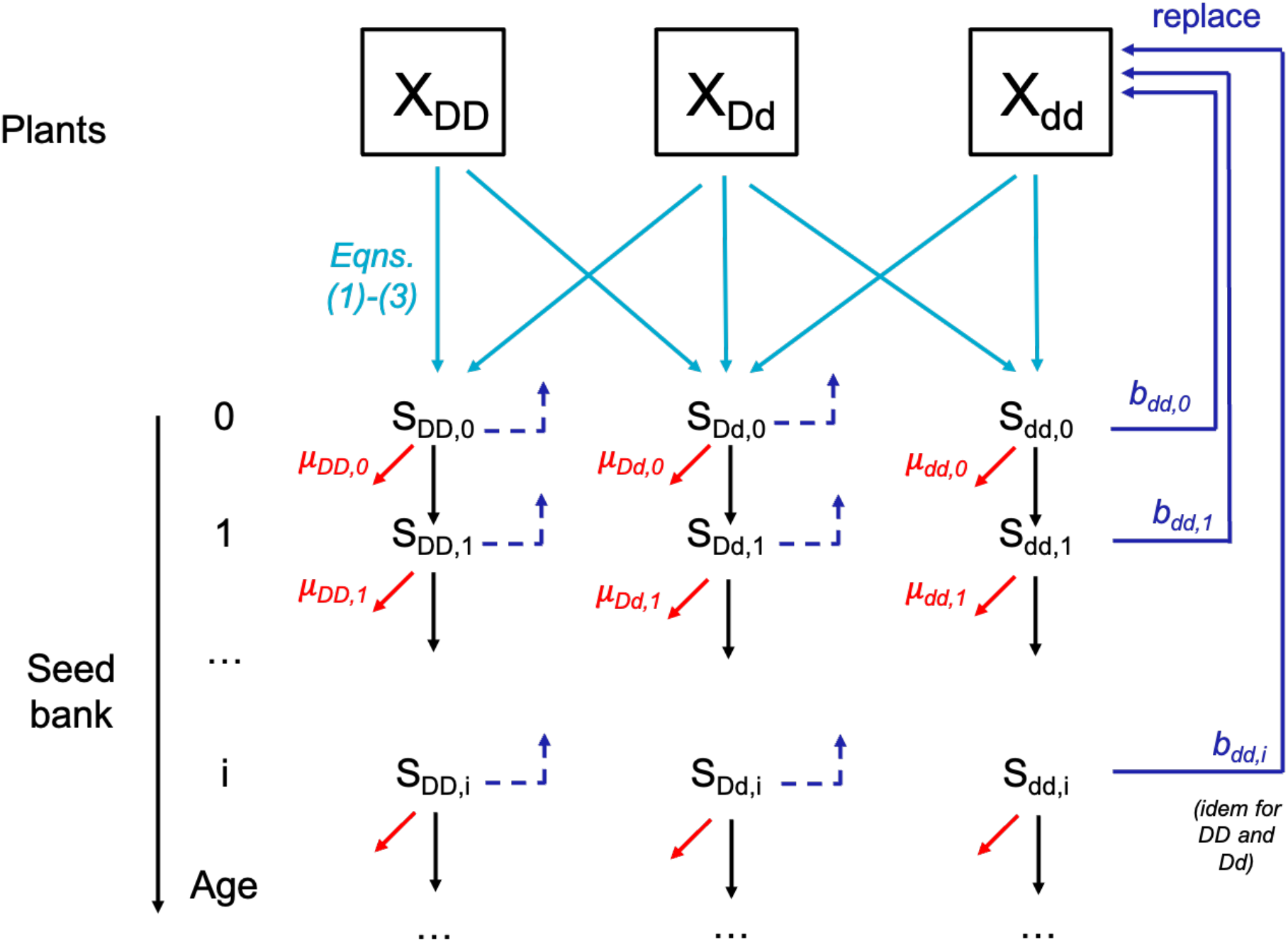
Seed bank model

**Supp. Figure 2.**
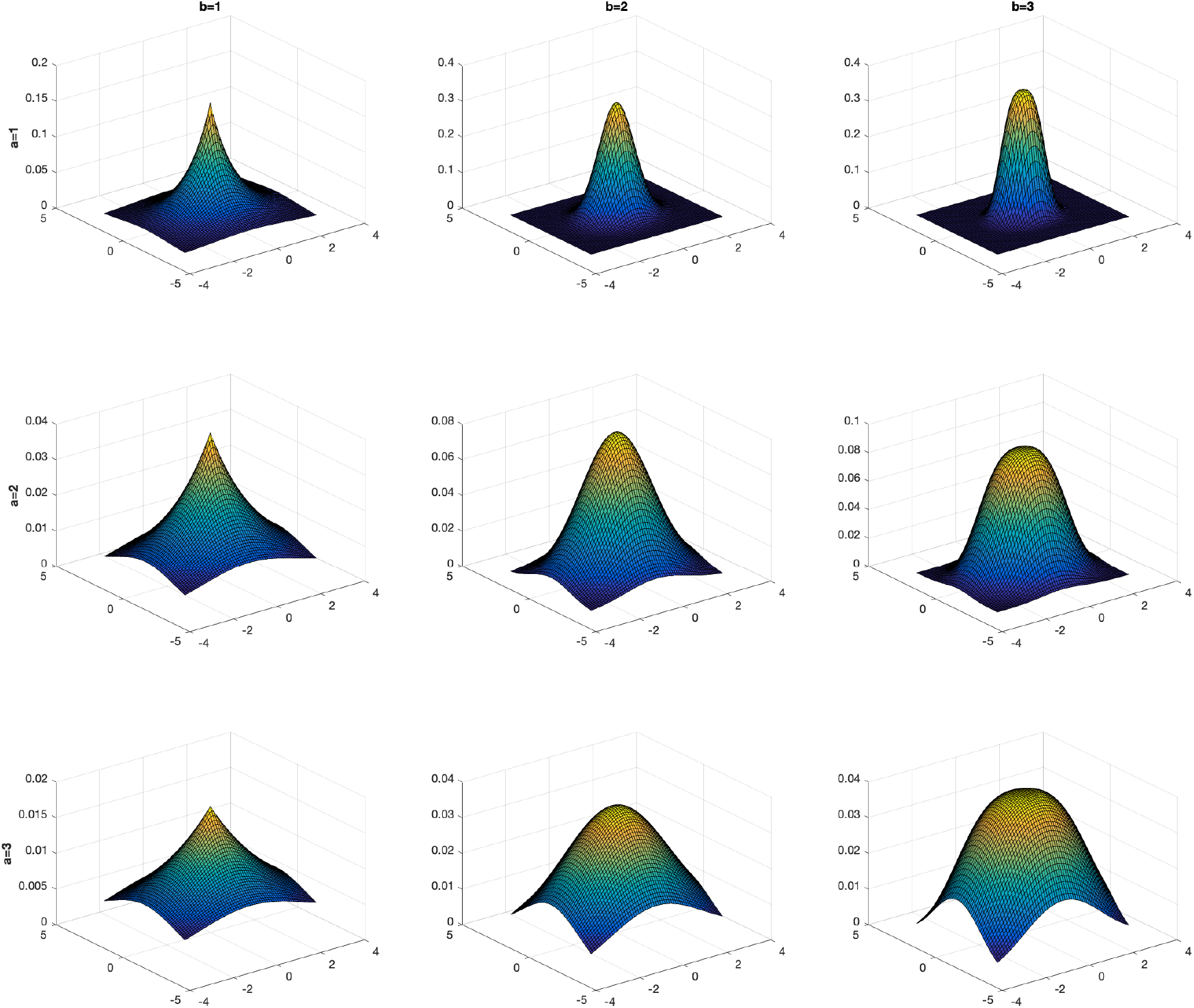
Dispersal kernels.

